# Falcon gut microbiome is shaped by diet and enriched in *Salmonella*

**DOI:** 10.1101/2022.11.25.517295

**Authors:** Anique R. Ahmad, Samuel Ridgeway, Ahmed A. Shibl, Youssef Idaghdour, Aashish R Jha

## Abstract

The gut microbiome is increasingly being appreciated as a master regulator of animal health. However, most avian gut microbiome studies have focused on birds of economic importance while the gut microbiomes of raptors remain underexplored. Here we examine the gut microbiota of 29 samples from four *Falco* species including hybrid birds— raptors of historic importance —in the context of avian evolution by sequencing the 16S rDNA V4 region. Our results reveal that evolutionary histories and diet are significantly associated with avian gut microbiota in general, whereas diet plays a major role in shaping the falcon gut microbiota. Multiple analyses revealed that gut microbial diversity, composition, and abundance of key diet-discriminating bacterial genera in the falcon gut closely resemble those of carnivorous raptors rather than those of their closest phylogenetic relatives. Furthermore, the falcon microbiota is dominated by Firmicutes and consists of *Salmonella* at appreciable levels. *Salmonella* presence may potentially alter the functional capacity of the falcon gut microbiota as its abundance is associated with depletion of multiple predicted metabolic pathways involved in protein mass buildup, muscle maintenance, and enrichment of antimicrobial compound degradation, thus increasing the pathogenic potential of the falcon gut and presents a potential risk to human health.

**Author Summary in Arabic:** 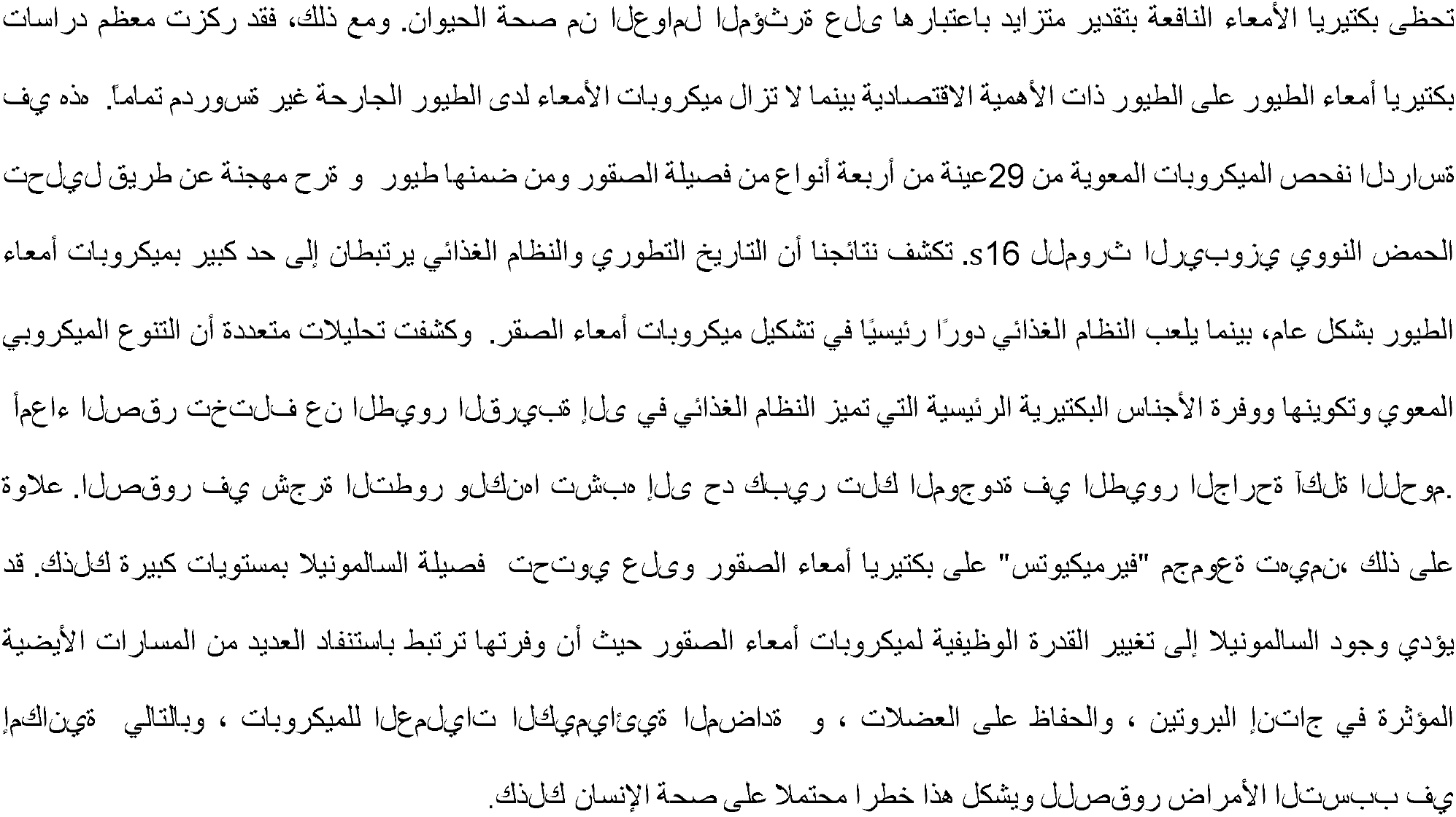

## Introduction

Animals harbor complex collections of symbiotic microorganisms including bacteria, archaea, viruses, and fungi that are collectively known as the microbiota (1). Emerging evidence indicates that these microbes provide their hosts with additional functions that animals have not yet been able to evolve, allowing them to fill novel ecological niches. Particularly important are the microbes residing in the gut as they can potentially regulate animal immunity, development, metabolism, behavior, and overall health (2-7). Because of the important role the gut microbes may play in maintaining animal health, it is essential to identify factors that contribute towards shaping the gut microbiota. The mammalian gut microbiota is strongly influenced by host evolutionary histories, diet, and environment (8-11) but how these factors contribute to the gut microbiome in birds remains unresolved (12, 13). Over 10 000 species of birds are distributed worldwide and they exhibit pronounced diversity in morphological traits, physiological functions, and adaptations to ecological niches (14). Consequently, the bird gut microbiota harbors hundreds of bacterial species belonging to diverse phyla that are associated with body size, flight, and dietary adaptations (15), indicating these bacteria play an important role in regulating physiological functions in birds (16). However, bird microbiome studies have predominantly focused on birds of commercial importance (17). Uber-carnivorous raptors whose diet almost exclusively consists of animal flesh, are highly underrepresented in the microbiome literature (18), with the exception of a few studies that include a handful of hawks, eagles, vultures, and falcons (15, 18-21).

Among raptors, *al suqur*—falcons of the genus *Falco* in Arabic—hold a special place in human history (22). Based on archeological records, *al suqur* have been connected to Middle-Eastern and Arab culture for the last 3 000 years (23). Falconry– the art of using falcons to hunt birds and small mammals –was practiced in Mesopotamia around 1 300 BC and falcons were one of the most frequently used hieroglyphic symbols in ancient Egypt (23). From Arabia, falconry spread via cultural diffusion to Central Asia and Northern Africa from where it spread to South Asia and Europe respectively in the Middle Ages (7th-11th Centuries) (24). Although falconry is currently practiced across the world (24), the Mesopotamian tradition of falconry is especially deeply ingrained in the Middle-East, where it is practiced as a traditional sport and *al suqur* are revered, beloved, and celebrated as cultural icons. Their cultural prominence is signified by their widespread mentions in Arabic literature and their use as emblems in coats of arms for several Middle-Eastern countries (22).

In addition to their cultural importance, birds of the genus *Falco* (henceforth referred to as falcons) are interesting from a biological perspective. They are a well-known raptors; yet, their phylogenetically closest relatives are Australaves that include passerines and parrots (25). Investigating the falcon gut microbiome can reveal several important insights regarding the role of gut microbiota in carnivorous birds. First, comparison of the bacterial species in the falcon gut with those of other birds may reveal the effects of diet and phylogeny on the avian gut microbiota. Second, it allows assessing the role of pathogens on the overall gut microbiota of birds as raptors are known to harbor pathogenic bacteria in their gut (26). Finally, publicly accessible raptor gut microbial data can be used in future conservation studies to evaluate the impact of human-induced environmental changes, which threaten to wipe out these endangered birds (12, 27).

Here, we generate and characterize the gut microbiomes of 29 falcons from four *Falco* species and hybrids using amplicon sequencing of the V4 region of the 16S rDNA gene. We integrate this dataset with a previously published dataset consisting of 636 birds (15) that includes representative species from each of the 9 clades spanning the entire avian phylogeny (25). Comparative analysis of the falcon gut microbiota in the context of avian evolution allows us to determine the role of their specialized uber-carnivorous diet in shaping their gut microbiota. Furthermore, we evaluate the contributions of *Salmonella*, a pathogenic bacterial genus, on the functional potential of the falcon gut microbiota.

## Materials and methods

### Ethics statement

This study was approved under an exempt protocol by the New York University IACUC Board. This observational study was carried out with consent from four veterinary clinics in the United Arab Emirates (Royal Shaheen Dubai, Al Sayad Falcons Abu Dhabi, SNC Falcons Abu Dhabi and Al Dhafra Abu Dhabi) who voluntarily donated fecal samples collected using non-invasive procedures.

### Sample and data collection

Fecal samples were collected from 29 captive *Falco* birds. Sampled species consisted of 11 purebred birds including *Falco cherrug* (n = 2), *Falco pelegrinoides* (n = 1), *Falco peregrinus* (n = 3), *Falco rusticolus* (n = 5), and 18 hybrid birds. Pure-bred and hybrid status was inferred from the records of the veterinary clinics. Freshly produced feces were collected with sterile inoculation loops and transported on ice-packs or dry ice to the laboratory within 4 hours of collection where they were stored at -80°C until DNA extraction.

### DNA Extraction

Bacterial DNA was extracted from the fecal samples using the Xpedition™ Soil/Fecal DNA MiniPrep kit (Zymo Research, Irvine, CA, USA) following the manufacturer’s protocol using roughly 0.25 g of fecal matter. Feces from the 29 birds were extracted in 1 to 4 replicates such that the total number of DNA samples was 42. DNA was eluted to a final volume of 100 μL and quantified on the Nanodrop 8000 (ThermoFisher Scientific, Waltham, MA, USA). The DNA concentrations and 260/280 ratios from the 42 fecal samples ranged from 6.9-107.7 ng/μL and 1.04-2.23, respectively.

### PCR and 16S rDNA gene amplicon sequencing

The V4 hypervariable region of the 16S rDNA gene was amplified using the 515F-806R primer combinations (28) with the FastStart High Fidelity PCR system (Roche Applied Science, Penzberg, Germany). Agarose gel electrophoresis (2% gel, 90 V, 400 mA, and 30 minutes running time) was performed to confirm successful amplification of DNA. PCR products were purified using AMPure XP beads, DNA concentrations of the purified products were measured using a Qubit (ThermoFisher Scientific) and molarity was calculated based on the size of DNA amplicons as determined by a 2100 Bioanalyzer instrument (Agilent Technologies, Santa Clara, CA, USA). The average amplicon size was 411 bp and the average concentration was 12.9 ng/μL (SD: +/-11.5). Samples were diluted to 2 nM based on library size as recommended in the Illumina 16S Sample Preparation Guide, pooled at equimolar quantities, and sequenced on the MiSeq platform using the Reagent Kit v3 (Illumina, San Diego, CA, USA).

### Computational analysis

A total of 6 058 432 single-end raw reads were obtained from the 41 falcon samples (one sample was dropped due to low read count). In order to analyze these reads, we developed an analyses workflow that was first tested using a previously published dataset by Song et al. (EBI accession number PRJEB35449) (15) that consisted of 16S rDNA V4 region forward reads generated using the same primers as our dataset (515F-806R) from 2,135 vertebrates, including 1 074 birds, 747 mammals, and 314 other animals.

Sequencing reads from the Song et al. dataset were processed with DADA2 (29) and analyzed using the phyloseq package (30) in R [4.2.0]. The reads were trimmed to 100 bp in order to retain high quality sequences (phred > 30). This dataset contained sequencing reads that were generated from 8 different runs on HiSeq and MiSeq instruments. Thus, sequencing data from each of these runs was processed individually. Reads with N nucleotides and/or > 2 expected errors were discarded (maxN = 0, maxEE = 2, truncQ = 2) and sequence variants were inferred from each sample. The falcon data from our sequencing run was processed with the same parameters and then merged into a single sequence table which resulted in a total of 272 103 907 reads. After chimera removal 261 167 457 (96%) reads were retained. Taxonomy was assigned using the RDP classifier (31) utilizing the SILVA v132 training set (32) as the reference. The R package DECIPHER (33) was used for multiple sequence alignment and a maximum likelihood tree was constructed with phangorn (34) using the neighbor-joining method.

A total of 255 109 025 reads (98%) belonged to the Song et al. (15) study, which consisted of 2 399 samples, including replicates. After retaining the replicates with highest coverage, a total of 229 672 486 reads belonging to 289 811 ASVs from 2 135 animals were initially retained. After removing ASVs with an abundance of less than 2 and presence in less than 5 samples, 187 673 522 reads and 24 267 ASVs remained from the 2 135 samples. Furthermore, non-bacterial reads (eukaryotes and fungi) and samples with sequencing depth of less than 10 000 reads were removed. These filtering steps resulted in a total of 187 014 663 reads (81%) and 24 267 ASVs from 1 824 animals. To prevent imbalances in species sample numbers, a maximum of 5 individuals per species were kept which resulted in a “comprehensive vertebrate dataset” (CVD) consisting of 136 976 389 reads and 24 065 ASVs from 1 330 animals from the Song et al. study, which was used to evaluate our workflow as described in Supplementary Figure 1.

Next, all non-avian samples were removed to create an “avian reference dataset” (ARD) which consisted of 61 469 984 reads and 10 603 ASVs from 675 birds spanning 9 phylogenetic clades (Supplementary File 1) (25). This dataset was used to assess the falcon microbiome in the context of broader avian evolution.

Finally, the falcon microbiome data was analyzed independently. This dataset consisted of a total of 6 058 432 reads belonging to 843 ASVs from 41 falcon samples (29 falcons and 12 replicate samples). After filtering with the parameters described above, 5 618 239 reads and 145 ASVs were retained. To determine if there were differences between replicates, alpha and beta diversity analyses were performed. Statistically significant differences were not observed between replicates (Supplementary Figure 2); thus, the samples with the highest coverage were retained resulting in an “falcon specific dataset” (FSD) consisting of 4 368 722 reads and 109 ASVs from 29 falcons.

### Gut microbial diversity analyses

Alpha diversity was measured using species richness and Shannon’s Diversity Index, calculated by rarefying reads to various depths between 1 000 and 10 000. One hundred iterations were performed at each depth and the mean values were used as the estimate of these measures in each sample. A maximum depth of 10 000 reads was chosen to include all individuals in the datasets (CVD, ARD and FSD). Kruskal–Wallis tests were used to assess the significance of differences in each of the alpha diversity metrics at the maximum depth (10 000 reads), by which the diversity had plateaued. Dunn’s post-hoc test was performed to assess pairwise differences between groups. Generalized linear mixed effect models were used to evaluate associations between both measures of alpha diversity (response variables) and metadata factors. For the ARD, phylogenetic clade, diet, flight status, GI tract region sampled, collection methods and captivity were used as explanatory variables. For the FSD four variables were available, namely age, sex, sampling location, and purebred status and all four were used as explanatory variables. The explanatory variables were treated to have fixed effects and random effects were assigned to each individual. P-value < 0.05 was considered statistically significant.

### Gut microbial composition analyses

Beta diversity was assessed at the genus level using the unweighted and weighted UniFrac as well as the Bray–Curtis distances calculated by log transformation of the non-rarefied 16S rDNA count data (30). Principal Coordinate Analyses (PCoA) were performed using the phyloseq package (30) and visualized with the *ggplot2* package in R (35). PERMANOVA was performed using the *vegan* package (36) using 10 000 randomizations where P-value < 0.05 was considered statistically significant.

### Clustering

Partitioning around medoids (PAM) clustering was performed on the ARD using the *cluster* package (37). Individuals were grouped into multiple clusters (Ku=u2 to 14) based on the top seven principal coordinate axes obtained from the weighted UniFrac distances. Goodness of clustering was assessed using a “gap” statistic with 1,000 bootstrap replicates.

### Predictive functional abundances

PICRUSt2 was used to predict the functional contents of the falcon gut microbiome using the ASV counts (38). A PCoA was performed for the 29 falcon samples with Bray-Curtis distances using the predicted MetaCyc (39) pathway abundances as features.

### Differential abundances of genera and function

Differential abundance of bacterial genera between dietary groups and predicted MetaCyc pathways between falcons with high, low, and no *Salmonella* load (*Salmonella* reads >100, 1-99, and 0 respectively) was assessed using a negative binomial generalized linear model using the *differential expression analysis for sequence count data version 2* (*DESeq2*) package (40). Genera and predicted MetaCyc pathways with absolute log2 [fold change] > 2 and adjusted pu<u0.01 were considered significant. Multiple testing corrections were performed by computing FDRs using the Benjamini–Hochberg method.

### Random forests

All random forest classifiers were constructed using five-fold cross validation repeated three times with 500 trees using genera as predictors. The data was partitioned into training and validation sets as described below. The R-package *randomForest* (41) was used to build the trees and accuracy was used to select the optimal model. To train the classifier for phylogeny inferred clusters from the ARD dataset, 20 individuals per cluster were used for training and the remaining were used for testing. To train the classifier for dietary groups in the ARD (flesh-eaters, herbivores and piscivores) 10 individuals were used for training and 5 for testing. To train the phylogeny-diet classifier in the ARD, 11 individuals per group were used for training and the remaining were used for testing. We assessed the performance of the classifiers by generating area under the receiver operating characteristic curves (AUC) using the R-package *ROCR* (42) and the default *varImp* function in the *caret* package (43) was used to calculate the variable importance.

## Results

### Sample description

We sampled freshly produced feces from 29 falcons across 4 veterinary clinics in the United Arab Emirates. Birds from Royal Shaheen Dubai included barbary (*Falco pelegrinoides*, n = 1), sakers (*Falco cherrug*, n = 1), and hybrids (n = 2). Birds from Al Sayad Falcons Abu Dhabi included gyrfalcons (*Falco rusticolus*, n = 2), peregrine (*Falco peregrinus*, n = 1) and hybrids (n = 5) and those from Al Dhafra Falcons Abu Dhabi included peregrine (*Falco peregrinus*, n = 2), gyrfalcons (*Falco rusticolus*, n = 2), sakers (*Falco cherrug*, n = 1) and hybrids (n = 10). A single hybrid sample was obtained from SNC Falcons Abu Dhabi. A detailed description of the diet, flying activity, sex, and age of the birds in this study is provided in Supplementary Table 1.

### Gut microbiome variation across vertebrates

In order to analyze the falcon fecal samples, we created a 16S analysis workflow (Supplementary File 2) and tested its utility by reanalyzing a previously published dataset by Song et al. (15). This dataset contained sequencing reads from the V4 region of the 16S rDNA gene, generated using the same primer set used in this study. After quality control (see Methods, S1 Fig A-F), removal of low coverage samples (reads < 10 000) and balancing at the species level of the animals (n ≤ 5), we generated a “comprehensive vertebrate dataset” (CVD, Supplementary file 1) consisting of 1 330 vertebrates, including 636 birds, 449 mammals, and 115 reptiles. Analyses of this dataset using our workflow reproduced several findings from the original manuscript (15). We found that the gut microbiome compositions varied by vertebrate class (p = 0.001, PERMANOVA, S1 Fig G-H) and the gut microbiome compositions of bats resembled that of flighted birds (*p* = 0.238, *Dunn’s post hoc test*, S1 Fig H). Gut microbial diversity was further associated with flight status, GI tract region sampled and sample collection methods (*p* = 0.001 for all, *PERMANOVA*, Supplementary Table 2). Alpha diversity, species richness and Shannon’s Index, was lower in flighted birds and bats compared to non-flighted mammals respectively (*p* < 2.2e-16 for both, *Dunn’s post hoc test*, S1 Fig I). Gut bacterial diversity was also lower for migratory birds relative to non-migratory birds (*p* = 0.017, *Kruskal-Wallis test*, S1 Fig J). Species richness was strongly associated with body mass in both mammals and birds (R^2^ = 0.450 and 0.367 respectively, *p* < 2.2e-16 for both, S1 Fig K). In both flighted (R^2^ = 0.343, *p* < 2.2e-16) and non-flighted birds (R^2^ = 0.334, *p* = 0.017) richness scaled with body mass albeit less than with non-flighted mammals. Species richness did not scale with body mass in bats (S1 Fig L).

### Avian reference dataset and factors associated with bird microbiomes

Next, we sought to identify the factors contributing to gut microbial variation in birds. To do so, we created an “avian reference dataset” (ARD) by retaining only the birds, and then by keeping samples only from the Song et al. study, we analyzed the microbiome of 636 birds with 5 representatives per species from each of the 9 avian clades (25). This dataset is severely underrepresented in falconiformes, although it consists of a small number of raptors including eagles and vultures (N=15). A Principal Coordinate Analysis of the ARD using weighted UniFrac distances revealed that the avian gut microbiota composition is significantly associated with biological factors such as bird phylogeny (clade), captivity, flight status and the gastrointestinal tract region sampled (*p* = 0.001, 0.001, 0.004, 0.001 respectively, *PERMANOVA*), although technical factors such as sample collection methods also contributed appreciably (*p* = 0.001, *PERMANOVA*). A multivariable analysis that consisted of biological and technical variables revealed that the first Principal Coordinate axis (PCo1) was strongly associated only with phylogeny (*p* < 2.2e-16, Generalized linear model). Paleognathae, a clade consisting of old-world birds such as ostriches, rheas and kiwis and Australaves, a clade consisting of recently evolved passerines and parrots, comprised opposite ends of the PCo1 spectrum with the rest of the clades occupying intermediary positions (Fig 1A). On the other hand, PCo2 was associated with technical factors such as the gastrointestinal tract region sampled (feces vs intestine) and sample collection methods (ethanol vs freezing or RNALater) (*p* = 0.002, < 2.2e-16, and 0.004, respectively, *Generalized linear model*). These observations were consistent when using unweighted UniFrac and Bray-Curtis distances as well as when samples across the avian clades were balanced (maxN = 50 per clade) to account for the dominance of Australaves in the data (Fig 1B). Moreover, birds could be clustered into five groups based on the microbiome data alone (Fig 1 B,C). Consistent with the PCoA, the Paleognathae (old-world birds) and Australaves (parrots and passerines) aggregated into distinct clusters (Clusters 1 and 5 respectively) while the remainder of the birds were spread across the five clusters (Fig 1C). Furthermore, a random forest classifier was able to differentiate these five clusters with high accuracy (out of bag error = 19%, area under curve = 1, 1, 0.97, 0.99, and 0.95 respectively, S2 Fig A). Alpha diversity was also associated with phylogeny (*p* = 0.001, *Kruskal-Wallis test)* with Palaeognathae having the highest alpha diversity and Australaves having the lowest among the 9 clades (S2 Fig B). These analyses collectively indicate that host evolutionary histories play an important role in shaping the avian gut microbiota.

**Fig 1.**
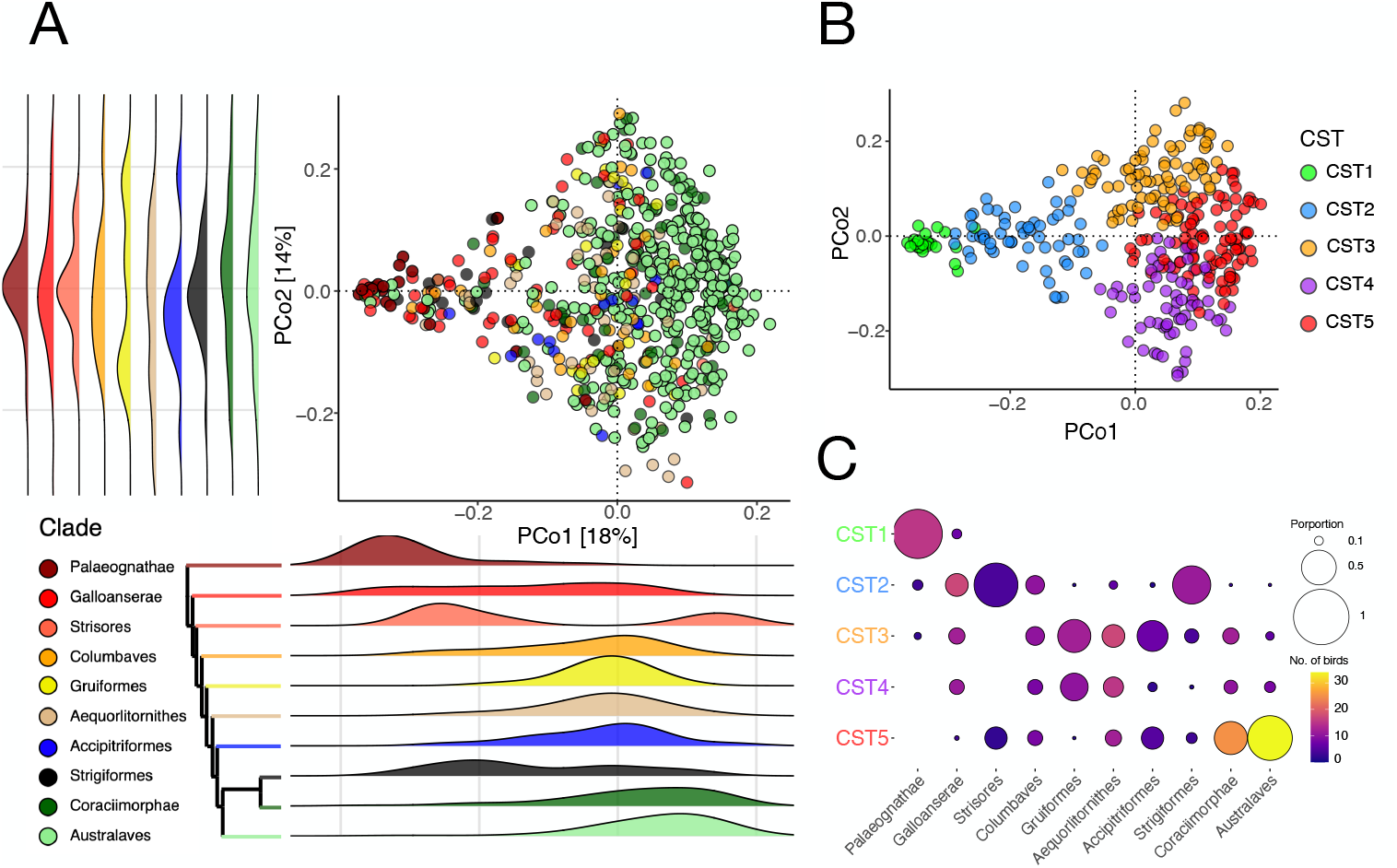
Bird microbiomes show signals of phylogeny. (A) PCoA of the weighted UniFrac distances of the avian 16S rDNA data colored by phylogenetic clades with each dot representing a bird. Palaeognathae (maroon) cluster on the left and Australaves (light-green) cluster on the right. Galloanserae to Coraciimorphae occupy transitory positions between Palaognathae and Australaves on PCo1 showing separation by phylogeny. (B) Clustering on balanced data (maxN = 50 per clade) from (A) suggests five major clusters. (C) Balloon plot showing number (color) and proportion (size) of clade distribution among the five clusters. Clusters 1 (green) and 5 (red) consist mainly of Palaeognathae and Australaves respectively.

Interestingly, neither the top PCo axes nor the clusters were clearly associated with diet, potentially reflecting the interspecies heterogeneity in diet and underrepresentation of flesh-eating carnivorous birds in this dataset (N=15). Thus, we hypothesized that a balanced dataset consisting of equal numbers of birds with distinct dietary habits may reveal the effect of diet on the bird gut microbiota. To evaluate this hypothesis, we identified a subset of birds from the intermediate clades (not Paleognathae and Australaves) whose diets were dominated by flesh, plants, and aquatic organisms. Flesh-eaters included uber-carnivorous raptors such as vultures and eagles (N=15), herbivores included hummingbirds and pigeons whose diet is dominated by plants and grains (N=15), and piscivores included penguins, flamingos and cranes that eat fish and crustaceans (N=15). PCoA using the weighted UniFrac distance on this balanced dataset revealed clustering by diet (R^2^ = 0.106, *p* = 0.001, *PERMANOVA*, Fig 2A). Moreover, a random forest classifier was able to assign the flesh-eating, herbivorous, and piscivorous birds to their respective source dietary groups with 80%, 100%, and 80% accuracies respectively (OOB error = 20% and AUCs = 0.96, 0.96 and 1, S2 Fig C).

**Fig 2.**
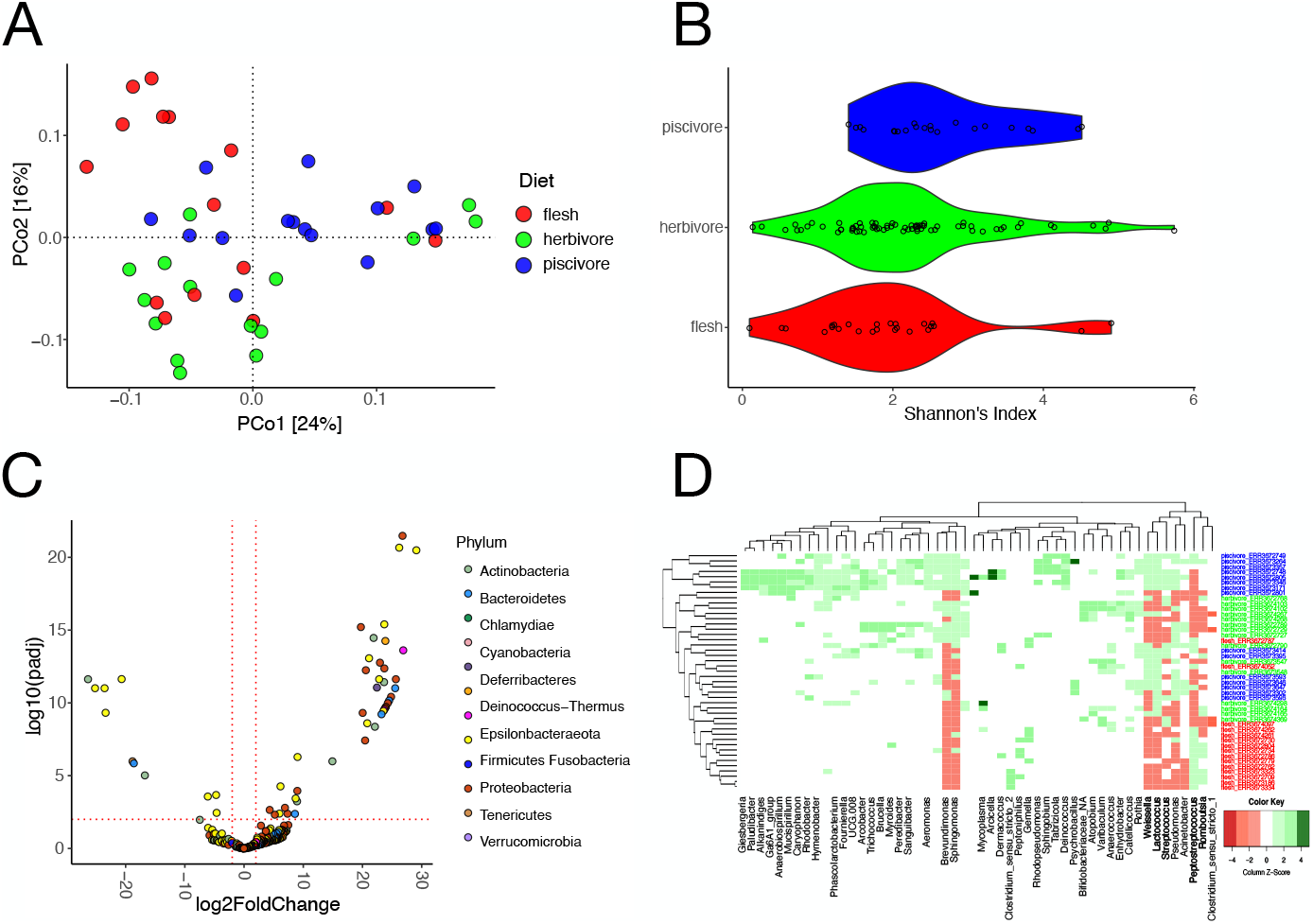
Bird gut microbiota variation by diet. (A) PCoA of birds with a flesh-eating diet (red), mainly vegetarian diet (green) and mainly fish-eating diet (blue). Diet accounts for 11% of the variation in the data (*p* = 0.001, *PERMANOVA*) in the weighted UniFrac ordination. (B) Shannon diversity is highest in piscivores (*p* = 0.01, 0.03 respectively) but not significantly different between flesh-eaters and herbivores (*p* = 0.099). (C) DESeq analysis reveals 52 bacterial taxa differently abundant between the dietary groups (log2fold > 2, *p* < 0.01). (D) Heatmap of the abundance of the 52 taxa from the DESeq analysis (x-axis), across the individuals of the three dietary groups (y-axis). The flesh-eaters separate from the other two groups.

Furthermore, alpha diversity also varied significantly between the three dietary groups (*p* = 0.002, *Kruskal-Wallis test*) (Fig 2B). Piscivores had the highest Shannon diversity but no significant differences were observed between the flesh-eaters and herbivores (*p* = 0.327, *Dunn’s post hoc test*). A total of 52 genera were differentially abundant in the flesh eaters compared to the other two dietary groups (log[fold change] > 2, *p* < 0.01, DESeq, Supplementary Table 3, Fig 2C). Flesh-eaters were enriched for *Peptostreptococcus* and *Romboutsia*, bacterial genera associated with high protein diets in animals (44, 45) and depleted in bacteria that assist in fiber digestion in the gastrointestinal tract namely, *Lactococcus, Streptococcus* and *Weissella* (46). Both *Peptostreptococcus* and *Romboutsia* were largely depleted in herbivores and piscivores (Fig 2D).

### The falcon microbiota resembles that of raptors

To assess the falcon microbiota in the context of bird evolutionary history, we processed the 16S rDNA V4 sequences generated from 41 samples sequenced from 29 falcons in this study using the same parameters used to generate the datasets analyzed above (described in Methods) (S3 Fig A-F). The replicate samples did not vary significantly from one another (*p* = 0.575, *PERMANOVA*, S3 Fig G,H). After removing the replicates by discarding the samples with lower coverages (see Methods), 4 368 722 reads from 29 falcons remained in the ARD. Principal coordinate analysis of this dataset using the weighted UniFrac distance reproduced the overall pattern observed in Fig 1 (Fig 3). The falcons from this study overlapped with the reference falcons in the ARD and they grouped with raptors (Fig 3A). Accipitriformes was the only clade, apart from the lowly sampled clade Strisores (n = 6), that was not significantly different from the falcons on both PCo axes (*p* > 0.05, *Dunn’s post hoc test*). This indicates that diet may have a significant effect on the falcon gut microbiota.

**Fig 3.**
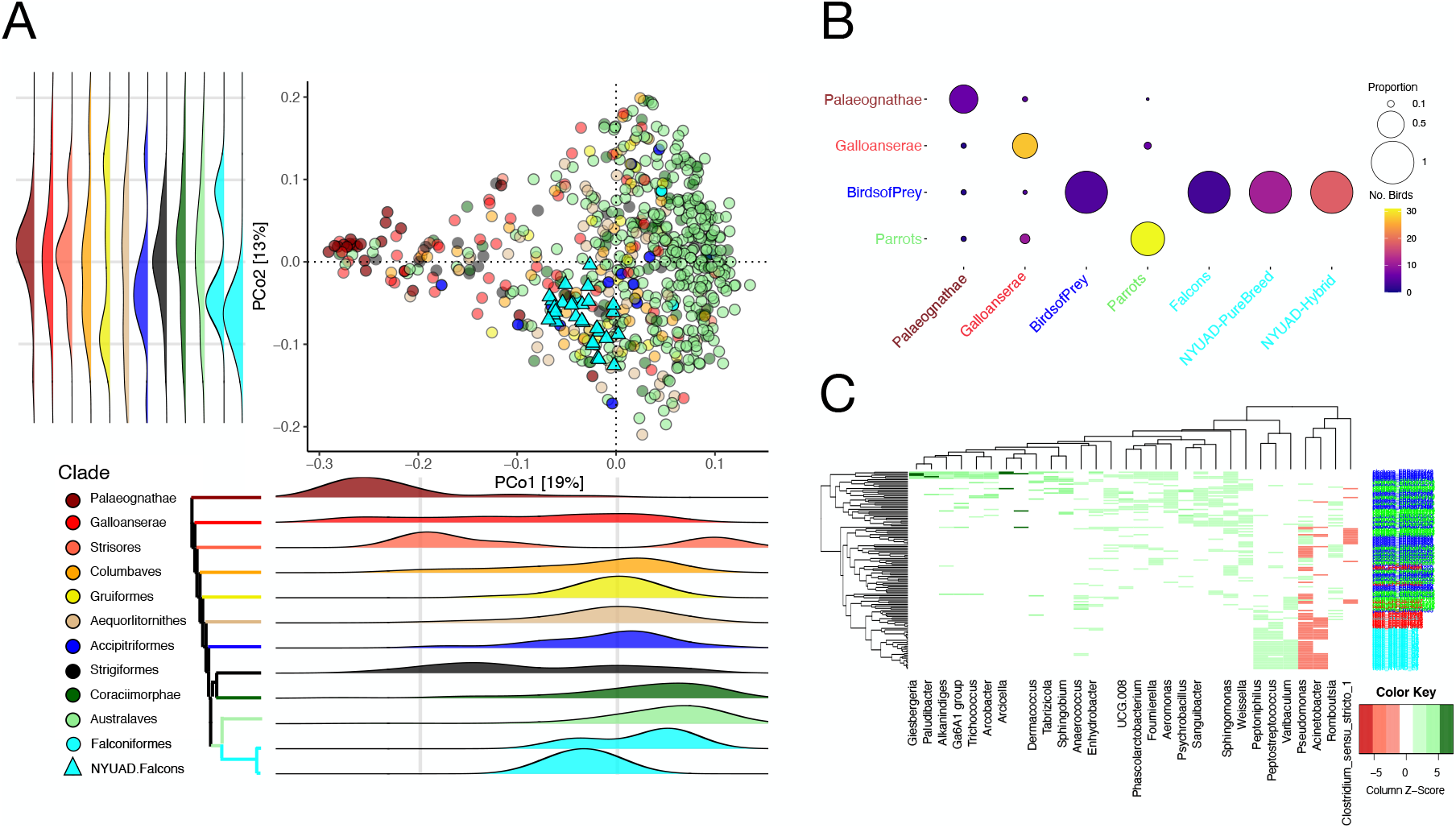
Falcon gut microbiota resemble those of raptors. (A) PCoA of the weighted UniFrac distances of the avian 16S rDNA data colored by phylogenetic clades with each dot representing an individual. Our falcons are shown as cyan triangles while falcons from the Song et al. dataset are shown as cyan circles. On PCo1 and PCo2 falcons and Accipitriformes (blue circles), all raptors, overlap with each other. (B) Balloon plot shows the accuracy of the random forest classifier testing dataset on the four groups and falcons from the reference dataset and our dataset with size of circle denoting proportion and color denoting number of individual birds. All three reference falcons get classified as raptors along with all 29 of our falcons showing that the falcon microbiome resembles that of raptors (common diet) and not parrots (genetically closer). (C) Heatmap of 30 taxa that are differentially abundant among the three previously identified dietary groups and also between our falcons vs herbivores and piscivores. Dendrogram from hierarchical clustering (left) separates individuals by a flesh-eating vs non-flesh-eating diet. The color red represents an underabundance of a genus and green represents an overabundance.

To further investigate the effect of diet on the falcon microbiome, we used the random forest model trained to classify birds into their dietary groups described above (S2 Fig C) and used 29 falcons from this study as the test dataset. All of the falcons from this study were classified as flesh-eaters. When parrots were used as the test dataset, 54% were grouped with herbivores and 44% with piscivores. We created a second random forest classifier model, this time encompassing both the phylogenetic and dietary groups to test if the falcon microbiomes resemble those of their phylogenetic relatives or those of raptors with whom they share dietary preferences. This model included 44 birds from the ARD (excluding falcons from this study) evenly distributed across four categories (N=11), three of which correspond to the clustering analysis above (Fig 1B): (i) old-world birds-Palaeognathae -representing Cluster 1, (ii) Galloanserae representing Clusters 2-4, (iii) parrots representing Australaves in Cluster 5 as well as the herbivores. For the fourth category, we included the raptors (Accipitriformes). If the falcon microbiota had stronger resonance with its phylogenetic cousins, they would be expected to cluster with parrots. If diet trumps phylogeny, they would cluster with the raptors. If their gut microbiota did not resemble either of these two groups, they would be expected to cluster with Galloanserae. The random forest model accurately classified the birds from the four source groups in the training dataset (OOB error = 11%). In the testing dataset, the model accurately classified 100% of the raptors (N = 4), 67% of the Palaeognathae (N = 9), and 79% of the parrots (N = 39) to their respective groups (Fig 3B). As expected, classification error was high for the Galloanserae because their gut microbiome does not cluster into a single group but spans multiple clusters (Fig 1C). All of the falcons in our dataset (N = 29) as well as the reference falcons included in the ARD (N = 3) were classified as raptors (Fig 3B).

Comparison of the 29 falcons in this study with the herbivores and piscivores included in the ARD revealed differential abundance of 85 genera (log[fold change] > 2, *p* < 0.01, DESeq). Out of these 85 taxa, 30 genera also discriminated against the three diet groups in the previous analyses including birds from the ARD (Fig 2C). Hierarchical clustering analysis using the relative abundance of these 30 genera revealed that the 29 falcons in this study cluster with flesh-eating raptors and not with the herbivores and piscivores in the ARD (Fig 3C). Both groups of flesh-eating birds are similarly enriched in *Peptostreptococcus* and depleted in *Weissella*. Neither of these genera were significantly different between the falcons in this study and raptors in the ARD (*p* = 0.09 and 0.47, *Kruskal Wallis test*). These analyses collectively indicate that the falcon gut microbiota resembles that of the raptors rather than that of their phylogenetically closest relatives— parrots —potentially owing to a large effect of carnivorous diet.

### *Salmonella* is associated with the falcon gut microbiome functions

We analyzed the gut microbiota of the falcons from this study independently of the avian reference dataset (Figure 4). Firmicutes was the dominant phylum (73% of total reads) in the falcon gut followed by Actinobacteria and Proteobacteria (Fig 4A). Gut microbiome composition did not vary significantly by sex, age or sampling site and no differences were observed between the purebred and hybrid birds sampled in this study (*p* > 0.05, PERMANOVA) (S3 Fig I). However, PCo1 obtained using weighted UniFrac distance was positively associated with species richness (Pearson’s r = 0.8, *p* = 0.001, *Generalized linear model*, Fig 4B).

**Fig 4.**
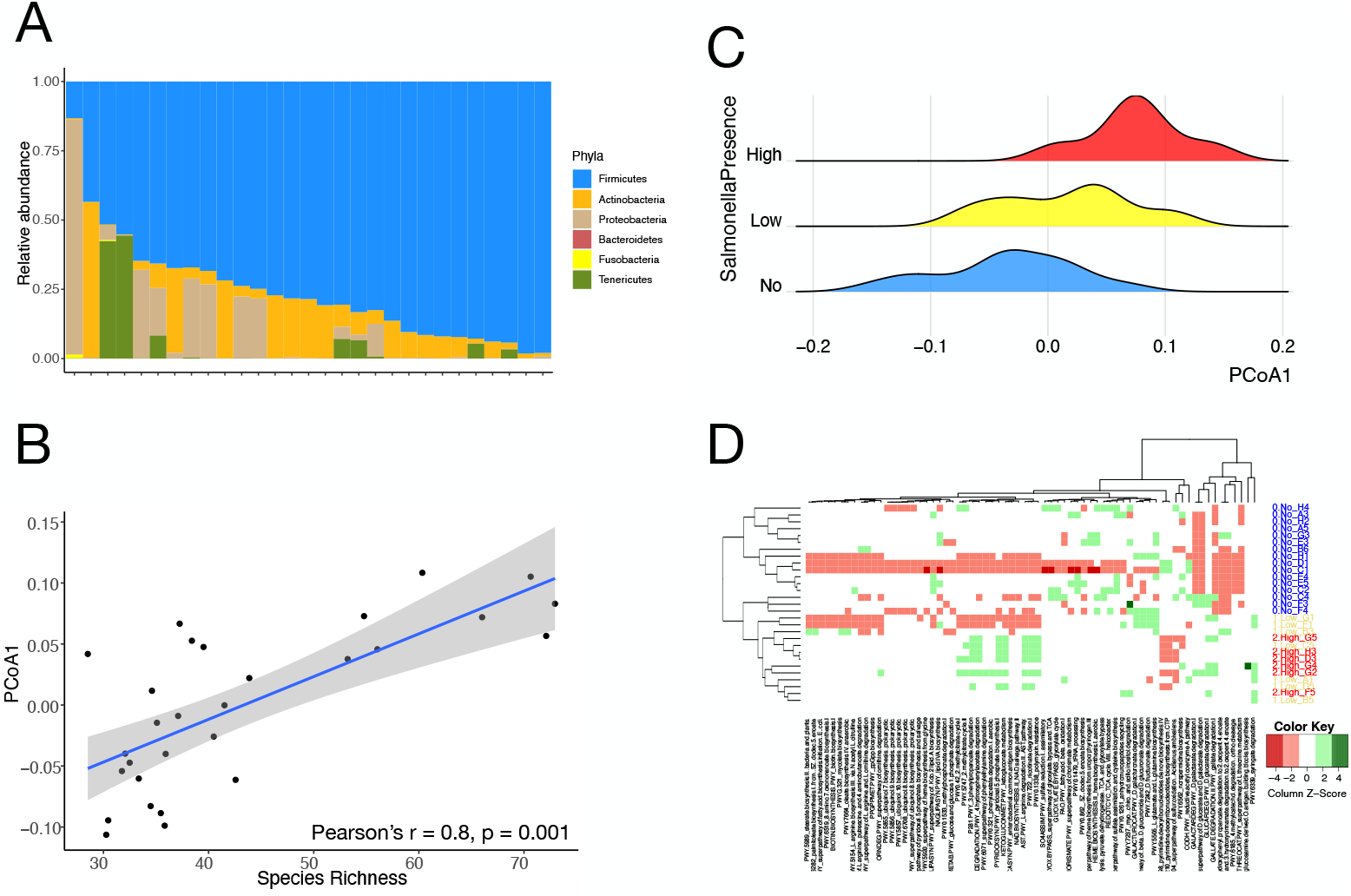
Falcon gut is dominated by firmicutes and Salmonella influences functional contents. (A) Abundance plot of phyla in the gut microbiome of 29 falcons, ranked by increasing firmicute dominance. Firmicutes is the most dominant phyla followed by Actinobacteria, Proteobacteria and Tenericutes. (B) PCo1 was strongly correlated with species richness (Pearson’s r = 0.8, *p* = 0.001). (C) PCoA analysis using Bray-Curtis distances of PICRUSt2-generated MetaCyc pathway aduncance data reveals difference in microbiome functional contents by presence of the bacterial genus *Salmonella*. PCo1 is strongly associated with *Salmonella* presence by three load levels (no, low and high) (D) Heatmap of 69 differentially abundant pathways calculated by a DESeq analysis. Dendrogram from hierarchical clustering (left) separates individuals by Salmonella presence (high and low loads vs no load) with differential abundance of several immunity and pathogenicity related pathways (top). The color red represents an underabundance of a pathway and green represents an overabundance.

We found *Salmonella* to be the 12^th^ most abundant genus in the falcons sampled in this study. All species of *Salmonella* are considered harmful pathogens for humans and they are naturally abundant in raptors (47). We assessed whether presence of *Salmonella* is associated with gut microbial diversity, composition, and functional potential of the falcon gut microbiota. Relative abundance of *Salmonella* was neither associated with the gut microbial diversity (*p* > 0.05, *Kruskal-Wallis test*) nor composition assessed using the first 3 PCo axes obtained using weighted UniFrac distances (*p* > 0.05, *Kruskal-Wallis test*). Next, we evaluated gut microbial functional potential using PICRUSt2 (38) to predict abundances of MetaCyc pathways. A Principal Coordinate Analysis using Bray-Curtis distance with the predicted pathways as features revealed the falcon gut microbial composition is strongly associated with *Salmonella* load (R^2^ = 0.2, *p* = 0.01, *PERMANOVA*) and not with age, sex, sampling site and hybrid status (*p* > 0.05, *PERMANOVA*). A clear differentiation along the PCo1 was observed between the falcons with high, low, and no *Salmonella* load (*p* = 0.003, *Kruskal-Wallis test*, Fig 4C). Furthermore, a total of 69 predicted pathways were differentially abundant between the falcons with no *Salmonella* compared to high or medium *Salmonella* loads (Fig 4D, Supplementary Table 4). Only a few of these pathways are specific to the *Salmonella* genome (n = 3). Several of these differentially abundant pathways are associated with protein buildup and maintenance, immune regulation, and breakdown of antimicrobial compounds, indicating that *Salmonella* presence may potentially alter the functional capacity of the falcon gut microbiota (Supplementary Table 5).

## Discussion

The modern birds radiated 66 million years ago (48) and today, there are over 10 000 species of birds that have adapted to diverse ecological niches worldwide resulting in tremendous morphological, physiological, and dietary specializations (49). Since the gut microbiome may play a critical role in regulating vertebrate physiology, it is important to understand its role in avian evolution and their adaptations across diverse habitats (12). Furthermore, understanding bird microbiomes is important for conservation efforts as human-activity-mediated changes in gut microbiomes of birds can be detrimental to their health and survivability (50). Finally, birds are known to harbor zoonotic pathogens in their intestines that can cause deadly human diseases (51). Despite its importance, the potential contribution of avian microbiome in bird evolution, conservation, and public health remains understudied because of several limitations. First, most bird microbiome studies have primarily focused on domesticated birds of economic value (17). Although microbiome studies of wild and captive birds using 16S rDNA sequencing are emerging, each study consists of limited samples– potentially reflecting difficulties in collecting samples from non-commercial birds (52) –which may lead to inconsistent findings across studies. Importantly, it is difficult to combine data from different studies due to variations in experimental techniques such as sample collection methods, DNA extraction protocols, and the choice of 16S rDNA region for amplification (12, 53), which prevent meaningful comparisons between studies. Therefore, it is imperative that future avian studies minimize variations in technical factors to make multi-study comparisons possible. A recent study (15) ameliorated some of these challenges by analyzing amplicons of the 16S rDNA V4 region using the universal primers (515F-806R) from 1 074 birds across the avian phylogeny along with hundreds of other animals, although raptors were severely underrepresented. In this study, we processed and reanalyzed this dataset and integrated much needed novel gut microbiome data from 29 falcons to create a comprehensive “avian reference dataset” consisting of 675 birds spanning 9 phylogenetic clades. The analysis of the avian reference dataset using our workflow reproduced previous results. For example, avian gut microbiota was significantly associated with physiological features such as flight status and captivity (15, 54). Therefore, this dataset and the workflow may serve as an important resource for the avian microbiome community.

In addition to corroborating previous findings, our results revealed a robust link between the avian evolutionary history and the gut microbiota. Multiple analyses revealed stark differences in both gut bacterial diversity and composition between the two most phylogenetically diverged bird groups – Paleognathae (old-world birds) and Australaves (recently evolved passerines and parrots), indicating that avian gut microbiota is shaped partly by the host evolutionary history. This finding is consistent with previous observations that the degree of similarity between microbiomes resembles evolutionary histories in birds (54, 55). The lack of gut microbial differences between the intermediate clades could be a potential consequence of the limited taxonomic resolution provided by the 16S rDNA reads. Future studies incorporating shotgun whole metagenomics sequencing may reveal additional bacterial strains and their functions relevant in avian evolution.

Our results also demonstrate that diet is significantly associated with the bird gut microbiota. The effect of diet on gut microbiota was not detected when 636 birds in the avian reference dataset were considered together, likely due to the dietary heterogeneity within each clade (56). However, analysis of a selected subset of birds whose diets are dominated by flesh, plant, or aquatic organisms revealed that diet has a significant effect on bird gut bacteria. Of the 52 genera differentially abundant between the three dietary groups, bacterial genera such as *Peptostreptococcus* and *Romboutsia* that are associated with high protein diets in vertebrates (44, 45) were also enriched in raptors whose diet are dominated by animal flesh. *Peptostreptococcus* is a genus of bacteria that is part of the normal gut flora of vultures (45, 57). Previous studies have found elevated levels of lactic acid bacteria such as *Lactococcus, Streptococcus* and *Weissella* in herbivorous birds compared to flesh eating birds (55). Presence of such bacteria helps metabolism of plant-based fibers via fermentation in the intestine (46, 58, 59). These bacteria were significantly depleted in the falcons in this study as well as in raptors generally, indicating that diet plays an important role in shaping the bird microbiomes.

Among the birds, the raptors are unique as their diet comprises almost exclusively of animal flesh. However, microbiome studies investigating their gut microbiomes are scarce. As a result, whether genetics, evolutionary histories, and diet have a measurable effect on the raptor gut microbiomes has remained unclear (18-21). We sequenced purebred and hybrid falcons of multiple species in this study and found that the gut microbiome composition of the falcons did not differ between sex, age, or purebred status, indicating that genetic variations in the falcon genome may have minor contributions in shaping their gut microbiota. By combining the falcon gut microbiota sequenced in this study with the avian reference dataset, we were also able to evaluate the potential effect of evolutionary histories and diet on the falcon gut. Our results show that the gut bacteria of falcons are more similar to those of raptors rather than phylogenetically adjacent parrots, indicating that diet has played a prominent role in shaping the falcon gut microbiota. This finding is consistent with previous observations that diet and flight associated adaptations are strong drivers of gut microbiome variations compared to host phylogeny in modern birds (54).

The falcon gut microbiomes warrant additional attention because they are known to harbor enteric zoonotic pathogens such as avian influenza virus (60), West Nile virus (61), Newcastle disease virus (62) and *Aspergillus* molds (63, 64) that can cause deadly human diseases and can be transmitted to humans via the practice of falconry (22). We found appreciable levels of *Salmonella* in the falcon gut and its abundance was strongly associated with shifts in predicted functional potential of the falcon gut (Supplementary Table 4). *Salmonella* high samples in our study showed elevated abundances of L-Arginine and L-Threonine degradation pathways. Both of these are amino acids that are essential for building protein mass, sustaining proper protein balance, and maintaining immune homeostasis in animals (65, 66). These findings indicate that the presence of *Salmonella* in the gut may negatively influence the overall health of the falcons by affecting the capacity to build and maintain protein mass, which is essential for flighted birds that engage in long-distance migrations. Moreover, small aromatic compounds such as gallic acid, 3-phenylpropionic acid, nicotinic acid, 4-methylcatechol, 3-hydroxycinnamate and their derivatives are known antimicrobials (67-71) and are known to have *Salmonella* inhibitory effects in humans (72). The falcons in our study that harbored high levels of *Salmonella* showed enrichment of functional pathways that degrade these antimicrobial compounds. Pathways associated with resistance of antimicrobial drugs such as polymyxin were also enriched in the falcons with high *Salmonella* load. These results indicate that *Salmonella* presence lowers the immune capacity of the falcons, making them susceptible to harboring infectious agents that can be transmitted to humans via falconry (73). Thus, *Salmonella* presence in the falcon gut not only threatens their well-being but it also presents a potential risk to human health. It is noteworthy that the presence of *Salmonella* in the gut of falcons in this study could be a consequence of captivity. Future studies investigating the gut microbiomes of wild falcons may reveal additional insights on the effect of this pathogen on the fitness of falcons in their native habitat.

Falconry is still practiced worldwide and it is a deeply rooted tradition in the Middle East where the falcons are carefully chosen and kept as pets, akin to dogs and cats in the western world. In addition, thousands of tourists interact with the falcons while visiting the Middle East. Our results point to the necessity of *Salmonella* screening in the falcons to reduce the potential of *Salmonella* outbreaks. In addition to *Salmonella*, the falcons may harbor additional pathogenic bacteria, which may not be detected in this study due to limited taxonomic resolution of the 16S rDNA sequencing. Future metagenomic studies that can distinguish between pathogenic and non-pathogenic bacterial strains are needed to identify these pathogens. Identification of bacterial strains that can counteract pathogenic bacteria can lead to probiotic supplements that reduce *Salmonella* and safeguard both human and falcon health.

## Acknowledgements

We thank M. M. Dieng, R. Shraim and M. Vinu for their contributions to the laboratory work and bioinformatics expertise respectively. We thank the Center for Genomics and Systems Biology, NYUAD Core Bioinformatics and Technology Platforms for assistance with technical work. We thank Al Sayad Falcons, Al Dhafra Falcons, SNC Falcons, and Royal Shaheen Events for providing access to falcon samples and for their assistance with sampling. We thank S. Al Dhabari, P. Bergh and the falconry community for the hospitality and expertise. This work is funded by NYUAD Capstone funds to Samuel Ridgeway, NYUAD grants ADHPG AD318 to A. R Jha and grant AD105 to Y. Idaghdour.

## Author Contributions

Conceptualization: A.R.J and Y.I. Sampling and laboratory work: S.R., Y.I. Data analysis: A.R.A and A.R.J. Interpretation: A.R.A, A.A.S, and A.R.J. Funding Acquisition: A.R.J. and Y. I. Supervision: A.R.J. and Y.I. Writing – original draft: A.R.A., S.R., A.A.S. and A.R.J. Writing - reviewing and editing: A.R.A., A.R.J., Y.I with input from all authors.

## Data availability

A phyloseq object containing the 16S data and metadata as well as the analyses protocols used in this work are included in the Supplementary Data. Sequence data is available in the GenBank SRA archive (PRJNA903547).

## Supplementary Data

**S1 Fig.**
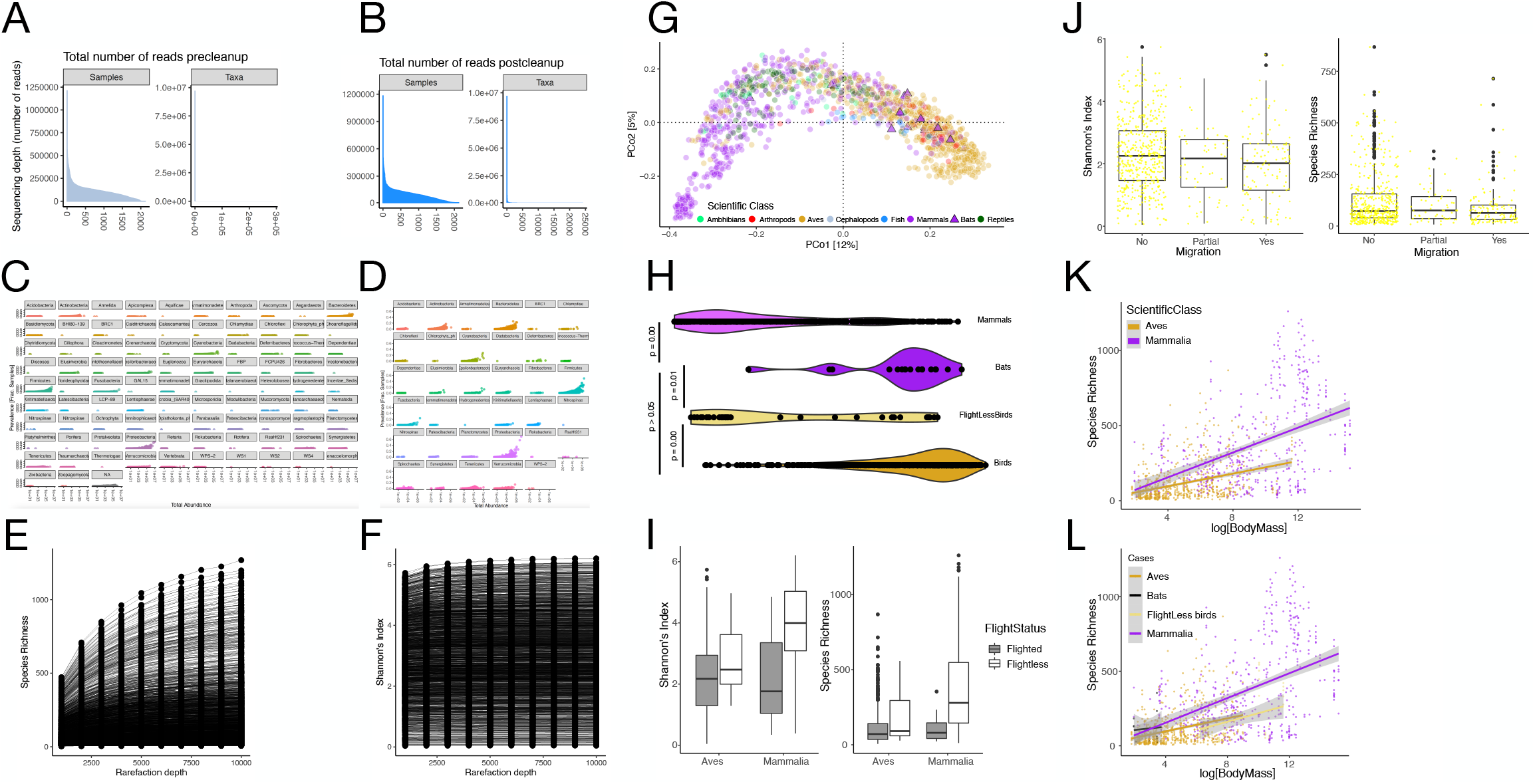
QC of Song et al. dataset and workflow validation. Sequencing depth for each sample and each taxon (A) before filtering any reads, and (B) After removal of lowly abundant ASVs; this does not reduce sequencing depth of either the samples or the taxa significantly. (C) Abundance of Phyla in the dataset. Fungal phyla were removed from the dataset. (D) After quality filtering, 35 phyla and a total of 24 000 taxa remained. Rarefaction curves for (E) species richness and (F) Shannon diversity show plateauing alpha diversity for most samples starting at a rarefaction depth of 10 000 reads. (G) Principal coordinate analysis of unweighted UniFrac distances at the ASV level and filtered to include a maximum of 5 individuals per species (1 330 points). Non-flighted mammals (purple dots) have a microbiome distinct from birds (yellow circles) and bats (purple boxes) cluster with birds. (H) Violin plot of PCo1 shows bird and bat microbiomes are not significantly different (*p* = 0.238, Dunn’s post hoc test). (I) Shannon diversity and species richness for flighted and non-flighted birds and mammals. Shannon for all pairwise comparisons had *p* < 0.05, except for flighted mammals vs flighted and flightless birds (*p* = 0.470, 0.111). For Richness all pairwise comparisons had *p* < 0.05, except for flighted mammals vs flighted and flightless birds (*p* = 0.361, 0.161). (J) Shannon for migratory and non-migratory birds have significantly different alpha diversities (*p* = 0.017). Richness between migration modes is not significantly different (*p* = 0.088). (K) Alpha diversity (richness) scaled against body mass for mammals (R^2^ = 0.450, *p* < 2.2e-16) and birds (R^2^ = 0.367, *p* < 2.2e-16). (L) In non-flighted mammals alpha diversity scales with body mass (R^2^ = 0.433, *p* < 2.2e-16) but in flighted mammals (bats) the relationship is insignificant (R^2^ = 0.0788, *p* = 0.808). In both flighted (R^2^ = 0.343, *p* = 0.018) and non-flighted birds (R^2^ = 0.334, *p* < 2.2e-16) alpha diversity scales with body mass albeit less than with non-flighted mammals.

**S2 Fig:**
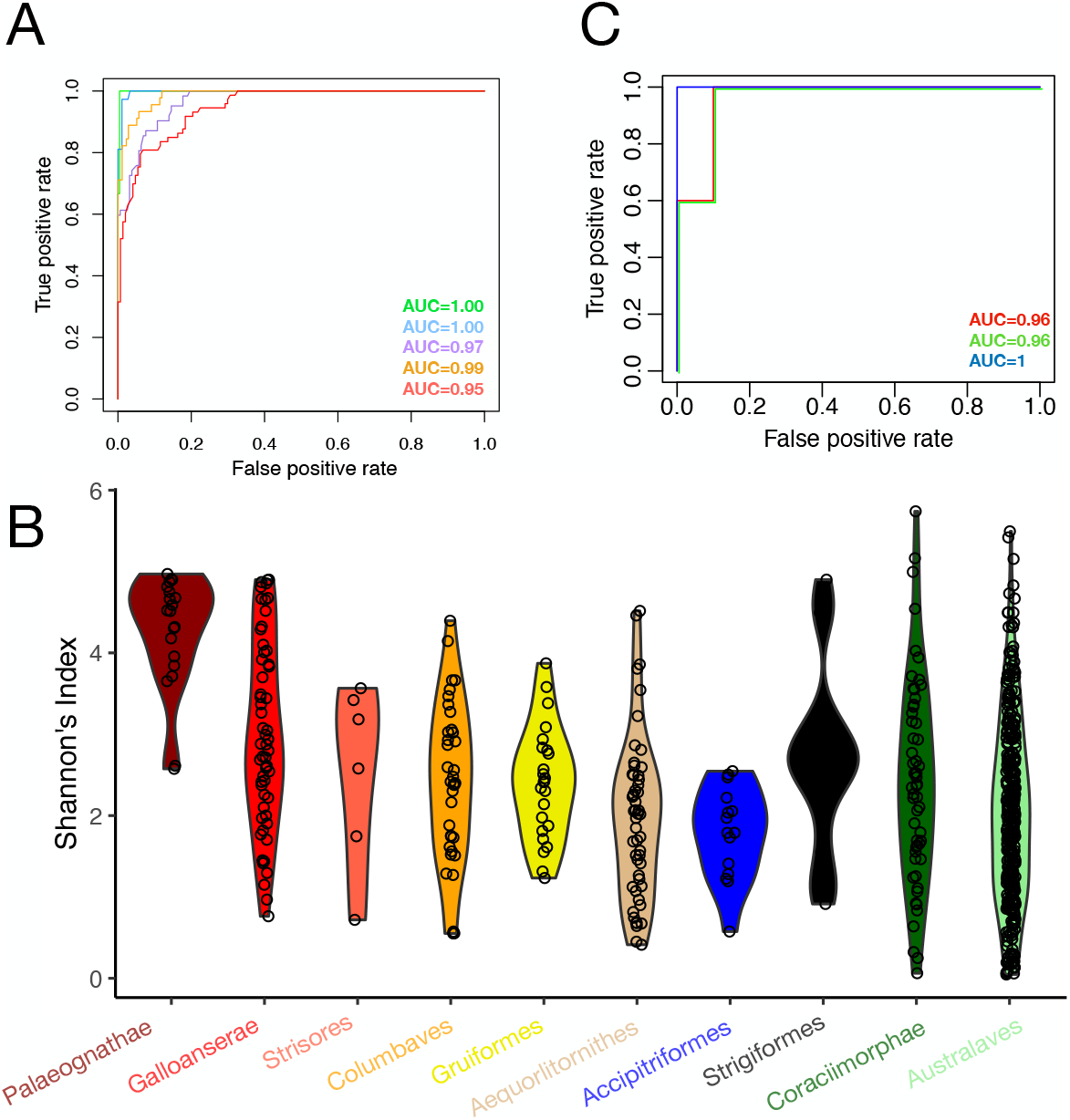
Phylogeny and diet supplementary figs. (A) A random forest classifier validates the grouping inferred from the clustering with 19% OOB error and AUCs = 1, 1, 0.97, 0.99 and 0.95. (B) Alpha diversity correlates with phylogeny, with Palaeograthae having the highest (p < 0.05 for all comparisons except with Galloanserae). (C) A random forest classifier supports the beta diversity analysis with 20% OOB error and AUCs = 0.96, 0.96 and 1 for the three dietary groups.

**S3 Fig.**
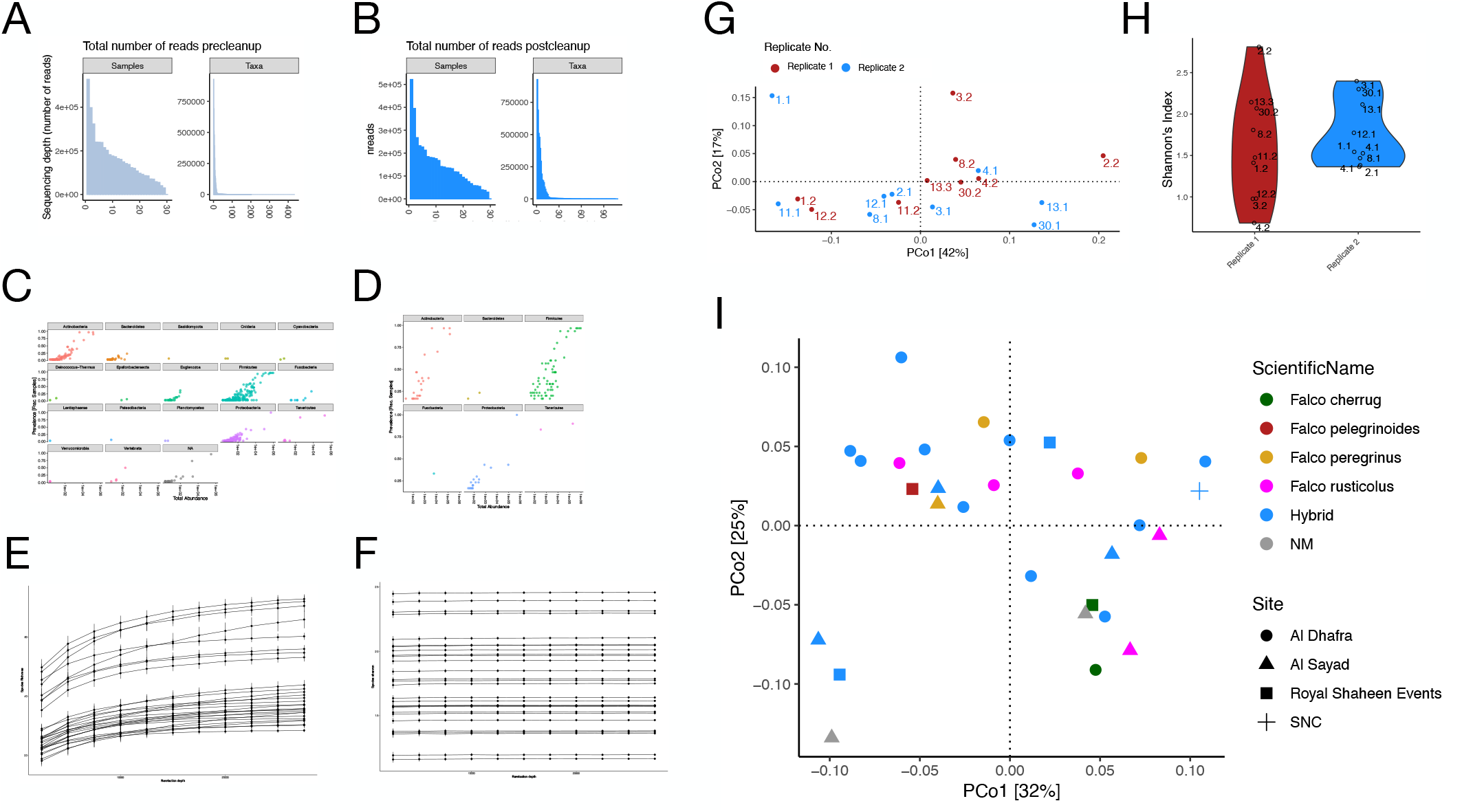
Application of workflow to falcon dataset and QC steps. Sequencing depth for each sample and each taxon (A) before filtering any reads, and (B) After removal of lowly abundant ASVs; this does not reduce sequencing depth of either the samples or the taxa significantly. (C) Total abundance and prevalence of phyla in the dataset. (D) After quality filtering, 6 phyla and a total of 109 taxa remained. One sample was removed on account of low reads, with the resulting phyloseq object having 29 samples. Rarefaction curves for (E) species richness and (F) Shannon diversity show plateauing alpha diversity for most samples starting at a rarefaction depth of 15 000 reads. (G) Beta diversity analysis of falcons with duplicates. No marked differences between replicates observed *p* > 0.05, PERMANOVA. Samples in blue are replicates with higher read counts and samples in red are replicates with lower read counts. (H) Alpha diversity between replicates does not differ markedly as well (*p* > 0.05 at rarefaction depth = 15 000). (I) PCoA of the weighted UniFrac distances of the falcon 16S rDNA data reveals separation that is not explained by species (color) or sampling site (shape).

**Supplementary File 1**

**Supplementary File 2**

